# RNA induces unique tau strains and stabilizes Alzheimer’s disease seeds

**DOI:** 10.1101/2022.01.29.478315

**Authors:** Amy N. Zwierzchowski-Zarate, Omar M. Kashmer, Josue E. Collazo-Lopez, Charles L. White, Marc I. Diamond

## Abstract

Tau aggregation causes neurodegenerative tauopathies, and trans-cellular propagation of tau assemblies of unique structure, i.e. strains, may underlie the diversity of these disorders. Polyanions have been reported to induce tau aggregation *in vitro*, but the precise trigger to convert tau from an inert to a seed-competent form in disease states is unknown. RNA triggers tau fibril formation *in vitro* and has been observed in association with neurofibrillary tangles in human brain. We tested whether RNA exerts sequence-specific effects on tau assembly and strain formation. Three RNA homopolymers, polyA, polyU, and polyC all bound tau, but only polyA triggered seed and fibril formation.

PolyA:tau seeds and fibrils were sensitive to RNase. The origin of RNA influenced the ability of tau to adopt a structure that would form stable strains. Human RNA potently induced tau seed formation and created tau conformations that preferentially formed stable strains in a HEK293T cell model, whereas other inducers produced strains that sectored. Finally, we found that soluble, but not insoluble, seeds from Alzheimer’s disease (AD) brain were sensitive to RNase. Thus, RNA specifically induces stable tau strains, and may trigger the formation of dominant pathological assemblies that propagate in AD, and possibly other tauopathies.

## Introduction

Tau forms highly ordered assemblies, termed amyloids, that underlie Alzheimer’s disease (AD) and related tauopathies [1], which may progress based on trans-cellular propagation [2] [3]. The fundamental origin of tauopathy is unknown, but systemic triggers include amyloid beta, trauma, and inflammation. Tau assemblies adopt faithfully self-replicating conformations in cells, termed strains. Strains may be propagated indefinitely in cell culture [4, 5], and induce specific patterns of neuropathology when inoculated into mouse models [6, 7]. We have observed that certain strains faithfully transmit neuropathology as prions [4]. However, so far, stable amplification and propagation of defined strains *in vitro* has proven difficult. It is unknown why tau forms unique strains, but the observation that a seed-competent tau monomer encodes limited strain ensembles[8], suggests that there may be very specific molecular triggers of different strains. Specific post-translational modifications of insoluble tau are reported to correlate with neuropathological diagnosis [9], but it is unknown whether phosphorylation triggers the conversion of inert to seed-competent tau monomer in tauopathy. Recent work from our lab has failed to find any detected post-translational modification of tau that correlates with this event [10].

Recombinant tau is thermostable and remarkably inert in solution. In the last ˜30 years, multiple inducers of tau fibril formation *in vitro* have been described[11], including polyanions such as heparin[12, 13], fatty acids[14], octadecyl sulfate[15], and RNA[16]. Tau was initially described to bind RNA, which sequestered it and prevented spontaneous tubulin assembly [17]. RNA has also been observed in association with tau tangles in brain samples[18, 19], and in association with induced tau aggregates in HEK293 cells[20]. Because tau has multiple positively charged residues, especially lysines, it was logical to assume that polyanions trigger tau assembly formation *in vitro* by neutralizing charge interactions and somehow unfolding the protein [21, 22] to facilitate its self-assembly.

The development of cell-based assays that amplify tau seeds, termed biosensors, has transformed our ability to characterize relatively small amounts of soluble tau assemblies[23, 24]. Biosensors are based on expression of full-length or repeat domain segments of tau (with or without disease-associated mutations) fused to suitable fluorescent protein tags. They respond to exogenous tau seeds by forming thioflavin positive inclusions[4]. Biosensor cell lines indefinitely and faithfully propagate myriad tau strains[4, 5], suggesting innate mechanisms of specific replication. Strains derived from cells will create specific, transmissible forms of neuropathology after inoculation into in a transgenic mouse model (PS19) that expresses tau (1N4R) containing a disease-associated mutation (P301S) [4, 6]. It is thus feasible to create and study tau strains induced *in vitro* and propagated in cultured cells. Given our interest in physiologic inducers of seed and strain formation, for which RNA seemed a plausible candidate, we investigated its role in this process.

## Results

### Tau binds polyA, polyC, and polyU RNA homopolymers

We first tested whether homopolymers of RNA would differentially bind recombinant tau. We used 40-mers of adenine (A), cytidine (C), and uracil (U), omitting guanidine because of its difficulty to synthesize. We purified recombinant, full-length (FL) tau monomer (2N4R) according to our prior methods[25], termed “tau “ henceforth. We immobilized tau on amine reactive second generation biosensors (ForteBio) and with exposure to increasing half-log concentrations of single-stranded RNA (0.13μM, 0.4μM, 1.3μM, 4μM, 13μM, 41μM), measured binding using biolayer interferometry (Octet, ForteBio). Binding data were interpolated according to a 2:1 interaction model. PolyA, polyU, and polyC strands each bound tau with similar avidities in the nanomolar range, as did a randomly generated single strand of DNA (Figure 1A; Supplemental Figure 1; Table 1).

**Table 1.** Binding of nucleic acids to tau. K_DAPP_ = Apparent binding affinity, k_on_ = association rate, k_off_ = dissociation rate. Results were calculated as the means of three independent experiments.

| Inducer | K<sub>DAPP</sub> | k<sub>on</sub> (M<sup>-1</sup> s<sup>-1</sup>) | k<sub>off</sub> (s<sup>-1</sup>) |
| --- | --- | --- | --- |
| polyA | 20 nM | 1.0 X 10<sup>4</sup> | 1.4 X 10<sup>-4</sup> |
| polyU | 254 nM | 1.7 X 10<sup>3</sup> | 4.2 X 10<sup>-4</sup> |
| polyC | 13 nM | 3.7 X 10<sup>4</sup> | 4.5 X 10<sup>-4</sup> |
| ssDNA seq1 | 12 nM | 2.8 X 10<sup>4</sup> | 3.4 X 10<sup>-4</sup> |

**Figure 1:**
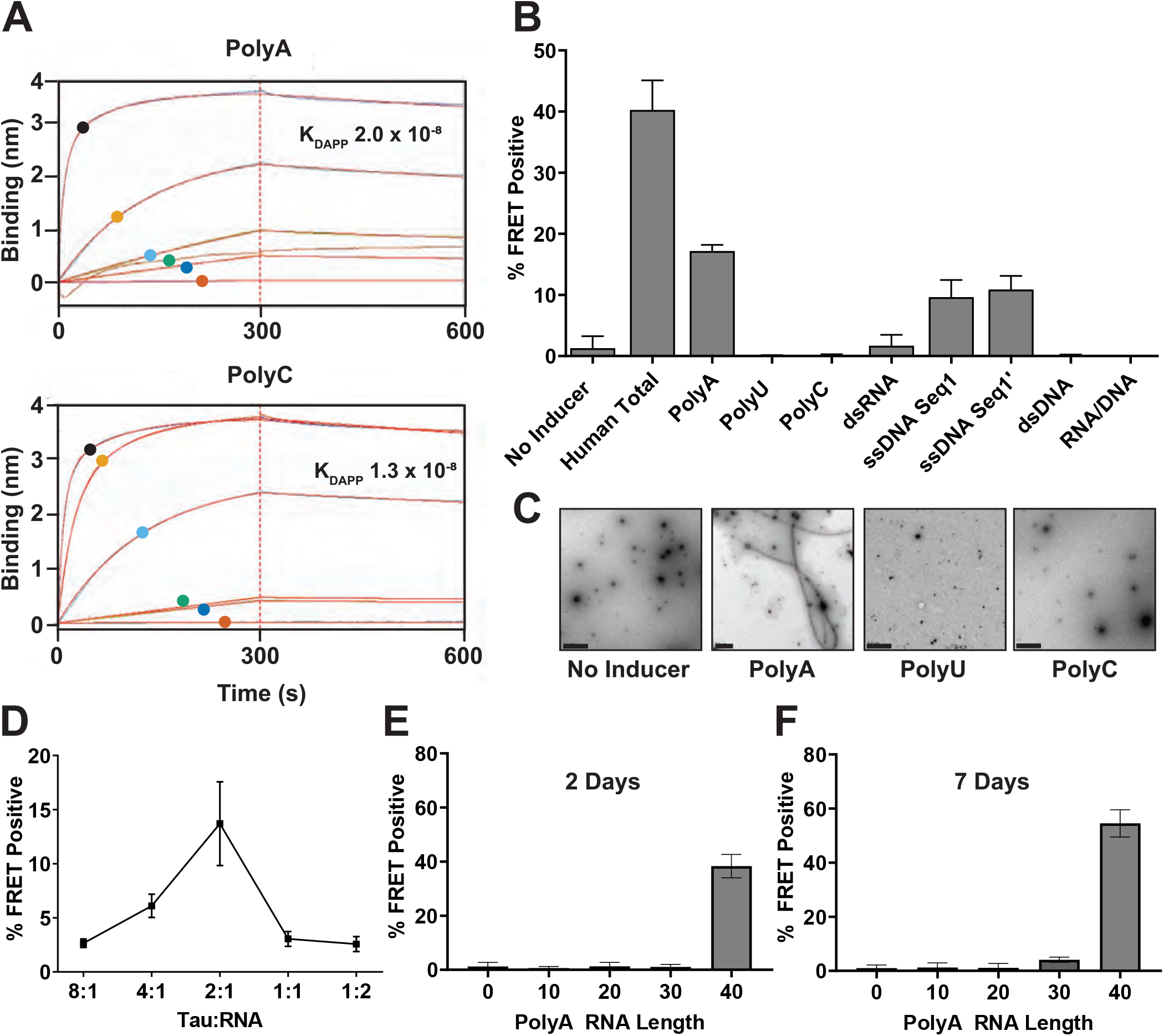
Tau binds multiple RNA polymers but only PolyA induces seeds. (A) BLI with tau immobilized onto AR2G biosensors and exposed to increasing concentrations of PolyA, or C RNA (A) BLI with tau immobilized onto AR2G biosensors and exposed to increasing concentrations of PolyU RNA, or ssDNA Seq1 (41μM 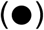,(13μM 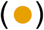,4μM 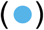, 1.3μM 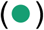, 0.4μM 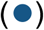, 0.13μM 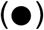). Tau bound all RNA with similar avidity, association, and dissociation rates. Figure shows representative data of PolyA or PolyC RNA binding to BLI AR2G sensors loaded with tau. K_DAPP_ calculated as mean of three independent experiments. Binding (nm) refers to the wavelength perturbation in reflected light from the biosensor. Red lines represent curves fit to primary data. See Table 1 and Supplemental Figure 1 for more data on these experiments. (B) Tau monomer incubated 24hrs with nucleic acid before transduction of v2L biosensors.Intracellular aggregation was quantified as % FRET positive via flow cytometry. Specific nucleic acids induced seed-competency in tau (Human Total RNA, PolyA RNA, ssDNA Seq1, ssDNA Seq1’) while others did not (PolyU RNA, PolyC RNA, dsRNA, dsDNA, RNA/DNA). (C) After 1 week with indicated inducer, tau was deposited on CF400 grids (4µM) and stained with 2% uranyl acetate. Grids were scanned with TEM for fibrils, and representative images are shown. No fibrils were seen for No Inducer, PolyU, or PolyC. Fibrils were observed only in conditions that induced seeding (PolyA). Scale bar =500nM. (D) Tau monomer incubated with various molar ratios of PolyA for 24 hr before induction of biosensors. 2:1 Tau to RNA (8µM:4µM) ratio produced optimal seeding. (E,F) Seeding of tau following incubation with various lengths (nt) of PolyA RNA for 2 (E) or 7 (F) days before transducing biosensors. >40nt was required for tau seeding. All seeding data represents two experimental replicates, each performed in technical triplicate. These experiments are representative of similar studies performed 19 times overall for (B), and five times overall for (D-F). Error bars = S.D.

### polyA induces tau seed formation *in vitro*

We next tested the ability of RNA homopolymers to induce tau fibril formation *in vitro*. We incubated tau (8µM) in the presence of polyA, polyU, and polyC RNA (50µg/mL, or 4µM) for 24 hours before measuring seeding with v2L biosensor cells that stably expressed the tau repeat domain with the P301S mutation fused to cerulean/mClover. After 48 hours, we quantified intracellular aggregation by fluorescence resonance energy transfer (FRET) as per prior studies [23]. PolyA generated seeding activity from tau monomer, while polyU and polyC did not (Figure 1B). A polyU/polyA hybrid also failed to elicit seeding. Two single stranded DNA homopolymers (polydA, polydT) did not induce seeding (Supplemental Figure 2A). Two randomly generated single-stranded, complementary DNA sequences of the same length and similar molecular weight as the RNA both effectively seeded, while the annealed double stranded complex did not, nor did an RNA/DNA complex of polyA hybridized with polydT (Figure 1B).

To rule out kinetic differences in seed induction, we incubated tau and RNA for up to two weeks. While polyA induced seed-competent forms, polyC and polyU never did (Supplemental Figure 2A-C). We observed fibril formation via transmission electron microscopy (TEM) that correlated with seeding activity after 48 hours incubation (Figure 1C). Thus, despite universal binding of RNA to tau, only certain single-stranded nucleotide sequences induced seed-competent conformations.

The optimal ratio of RNA:tau to induce seeding was 2:1 (Figure 1D). We defined the minimal length of RNA necessary to induce seeding by incubating tau with increasing sizes of polyA ranging from 10 to 40 nucleotides (nt) for various time periods. At 48 hours the smallest length of RNA capable of inducing seed-competent tau to a significant degree was 40 nt (P<0.0001, one-way ANOVA, post-hoc Dunnett’s multiple comparisons test) (Figure 1E). After 7 days we observed minimal increases in seed formation with 30 nt (4.1% FRET positive), which was still not significant compared to buffer control (P=0.1451, one-way ANOVA, post-hoc Dunnett’s multiple comparisons test), while 40 nt seeding increased from 38% to 55% FRET positive.

### RNA stabilizes seeds and fibrils

Amyloid fibrils are thought to occupy a particularly stable, low-energy state[26], which might predict their persistence after inducers are removed. We tested this idea for RNA induction. We first formed tau fibrils by incubation with polyA RNA. We then exposed fibrils to a mixture of RNase A and RNase T1 for 24 hours to degrade the RNA, and tested the effect on seeding and fibril integrity. Tau seeds induced by RNA lost all seeding activity after RNase exposure, while incubation with DNase or Heparinase had no effect (Figure 2A). To exclude a direct effect of RNase on biosensor activity, we added RNase directly to the seeds without pre-incubation, and detected no loss in seeding activity (Figure 2A). We next used TEM to test fibril stability in the presence of nucleases. RNase eliminated all detectable fibrils, but not DNase or Heparinase (Figure 2B). Conversely, fibrils induced by heparin remained unchanged after DNase and RNase treatment but disassembled and lost 74% of seeding after incubation with heparinase (Figure 2A,B). Incomplete loss of seeding after removal of heparin is consistent with our prior observations that it converts tau monomer to a stable, seed-competent conformation[27];[28].

### RNA influences strain composition

Tau adopts multiple, faithfully propagated assembly structures that produce unique, transmissible patterns of neuropathology *in vivo*, termed strains [4, 6]. Given that RNA sequence dictated seeding activity, we tested if it might also control strain composition. We previously developed the DS1 biosensor cell line, which expresses tau RD containing two disease-associated mutations (P301L and V337M) fused to YFP[4]. Although it represents a rarified experimental system, this line has proved very useful because it readily propagates myriad tau strains. Depending on their replication efficiency, some strains propagate faithfully, whereas others will sector, i.e. inclusions will steadily disappear from the cell population. Thus, one simple and robust classifier of strain identity is the ability to maintain itself in dividing cells or not.

Tau was incubated in the presence of total human RNA, total yeast RNA, polyA RNA, or ssDNA before treating DS1 cells with subsequent tau seeds to induce inclusion formation. After 72 hours, we used fluorescence activated cell sorting (FACS) to isolate 384 single aggregate-containing cells for each condition in the individual wells of 96-well plates (which was possible based simply on gating for the high fluorescence intensity that occurred when RD-YFP aggregates [8]). After 2 weeks of cell growth, we counted amplified colonies (n=130 derived from human RNA, n=228 from polyA, n=220 from DNA, n=98 from yeast RNA) that still propagated aggregates or had sectored. 60% of colonies induced by human RNA:tau and 43% of colonies induced by polyA:tau still contained inclusions, while only 1% of colonies induced by yeast RNA:tau and 3% of colonies induced by ss DNA:tau contained inclusions (Figure 3A). Hence, despite starting with 100% inclusion-bearing cells derived from tau seeds across multiple inducers, human RNA most efficiently induced conformations of tau that created stable strains.

**Figure 2:**
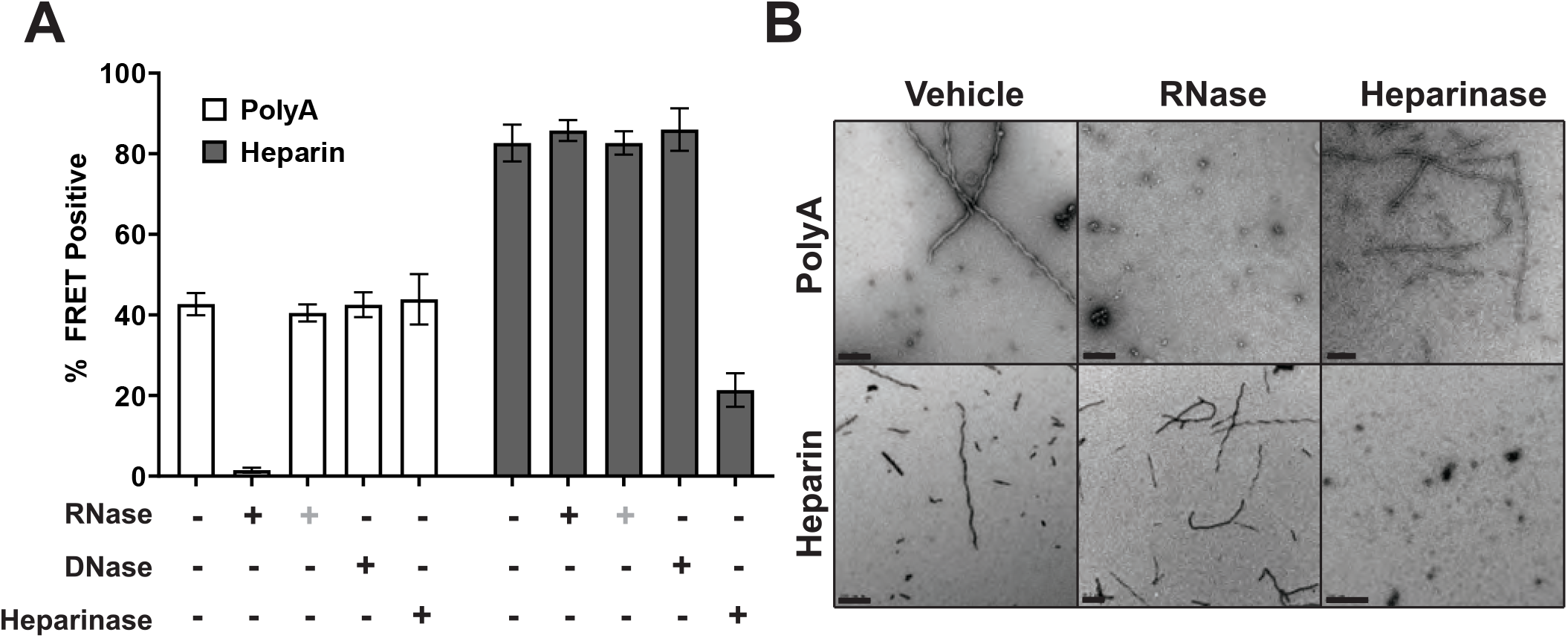
Stabilization of tau fibrils and seeds *in vitro*. (A) Tau fibrils pre-formed with PolyA RNA or heparin were treated with RNase (T1 and A, 316 U/μL and 3.2 mg/mL), DNase (I, 633 U/μL), or heparinase I (3797 U/mL) for 24 h before transduction of biosensors. (**+**) signifies RNase treatment added to fibrils at same time as seeding, with no pre-incubation. Seeds formed with PolyA were sensitive to RNase treatment.RNase added directly to seeds at the time of seeding had no effect. DNase and heparinase treatment did not affect seeds formed by PolyA. Seeds formed with heparin were sensitive to heparinase treatment, but not RNase or DNase. Seeding data represents two experimental replicates, each performed in technical triplicate. These experiments are representative of similar studies performed 15 times overall. Error bars= S.D. (B) TEM 400CF grids containing tau (4μM) stained with 2% UA. All grids were scanned; representative images are shown. Scale bar = 200nM. Fibrils formed by polyA lose integrity after exposure to RNase, but not Heparinase. Heparin fibrils were undetectable after heparinase treatment, but not RNase.

**Figure 3:**
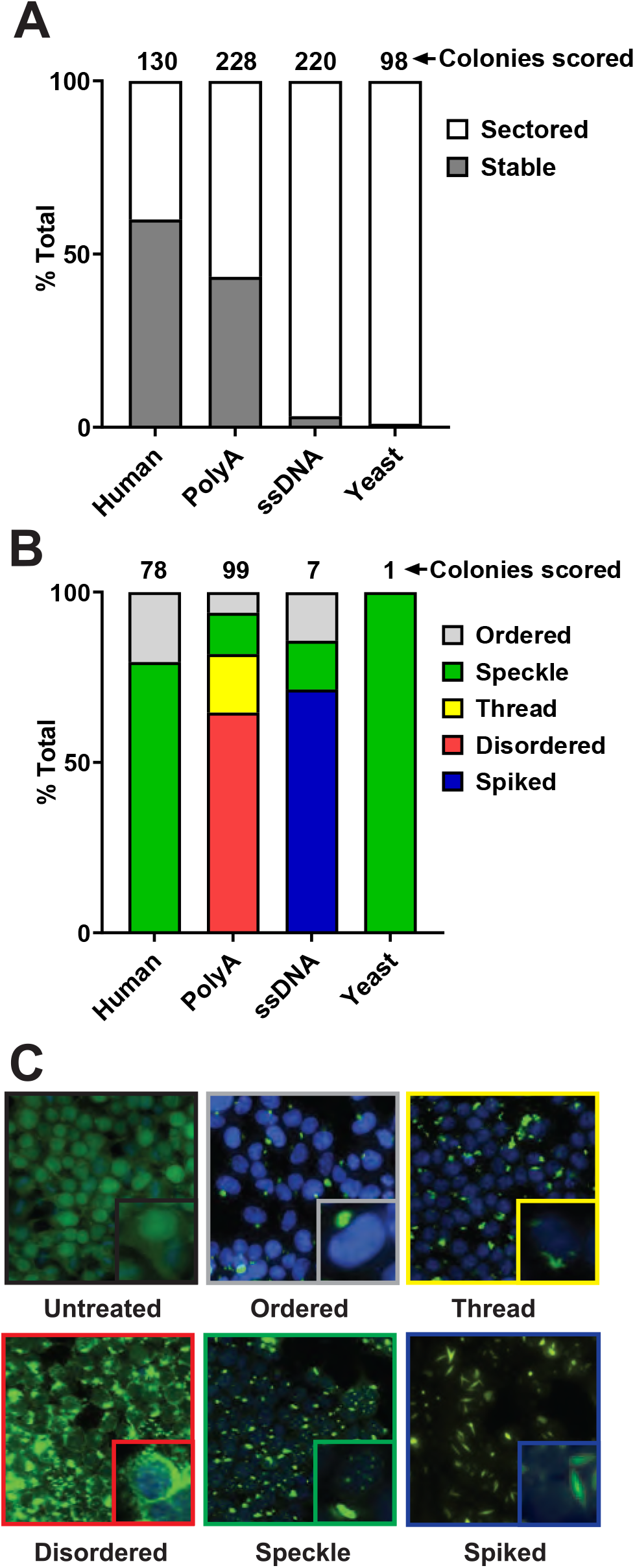
RNA inducer dictates aggregate maintenance and strain identity. (A) Tau seeds derived from different nucleic acid sources were transduced into DS1 cells and single cells were sorted into 96-well plates and monitored over time. Certain nucleic acids (Human, PolyA) created more stable strains than others (ssDNA, yeast), which lost aggregates, or sectored almost completely. N=130 derived from human RNA, 228 from polyA, 220 from DNA, 98 from yeast RNA. (B) Stable colonies containing inclusions were imaged using an In Cell Analyzer 6000 and resulting images were scored for morphology, blinded to the method of seed induction. Distribution of morphologies varied by inducer. N=78 derived from human RNA, 99 from PolyA, 7 from ssDNA, 1 from yeast. (C) Exemplars of morphologies.

To further explore the effect of RNA origin on strain identity, we maintained remaining stable colonies (n=78 human RNA, 99 polyA RNA, 7 ssDNA, 1 yeast RNA) for characterization of inclusion morphology, a surrogate for strain identity [4]. Colonies were assessed via fluorescence microscopy, and blindly scored for morphology. After unblinding and quantification, human RNA derived colonies largely had speckle morphology (79%), while remaining colonies were ordered (21%) (Figure 3B,C).Colonies derived from polyA RNA were primarily disordered (65%), in addition to thread (17%), speckle (12%), and ordered (6%) morphologies (Figure 3B,C). While relatively fewer ssDNA colonies stably maintained aggregates, we observed a distinct spiked morphology in 71% of colonies, and additional colonies with speckle (14.5%) and ordered (14.5%) patterns (Figure 3B,C). The sole remaining yeast RNA-derived colony produced speckle morphology (Figure 3B,C). Although distinct inducers of intracellular aggregates generally displayed a heterogenous profile of intracellular morphologies, the predominant morphology for each inducer was unique.

### Soluble AD seeds are stabilized by RNA

Given the ability of RNA to facilitate tau seeding *in vitro* and strains in cells, we tested its stabilization of seeds derived from the most common tauopathy, Alzheimer’s Disease (AD) [29]. We homogenized frontal lobe tissue from an AD patient and fractionated the tau based on sarkosyl-insoluble material, which produced fibrils (Supplemental Figure 3) and soluble fractions, using size exclusion chromatography to resolve soluble assemblies of different sizes. Because each sample had different amounts of seeding activity, to ensure that we evaluated seeding activity accurately across samples we first titrated each to empirically determine amounts in the middle range of a dose-response curve (Supplemental Figure 4). We then exposed each fraction to RNase, DNase, or buffer. After 24 h, we quantified seeding activity using biosensor cells. In total and soluble fractions, we observed a ˜70-85% reduction in seeding after RNase treatment (Figure 4). By contrast, fibrillar tau exhibited no change in seeding in presence of RNase. DNase had no effect on seeding for any fraction. These data indicated that RNA maintains soluble, but not insoluble, tau seeding in AD brain.

**Figure 4:**
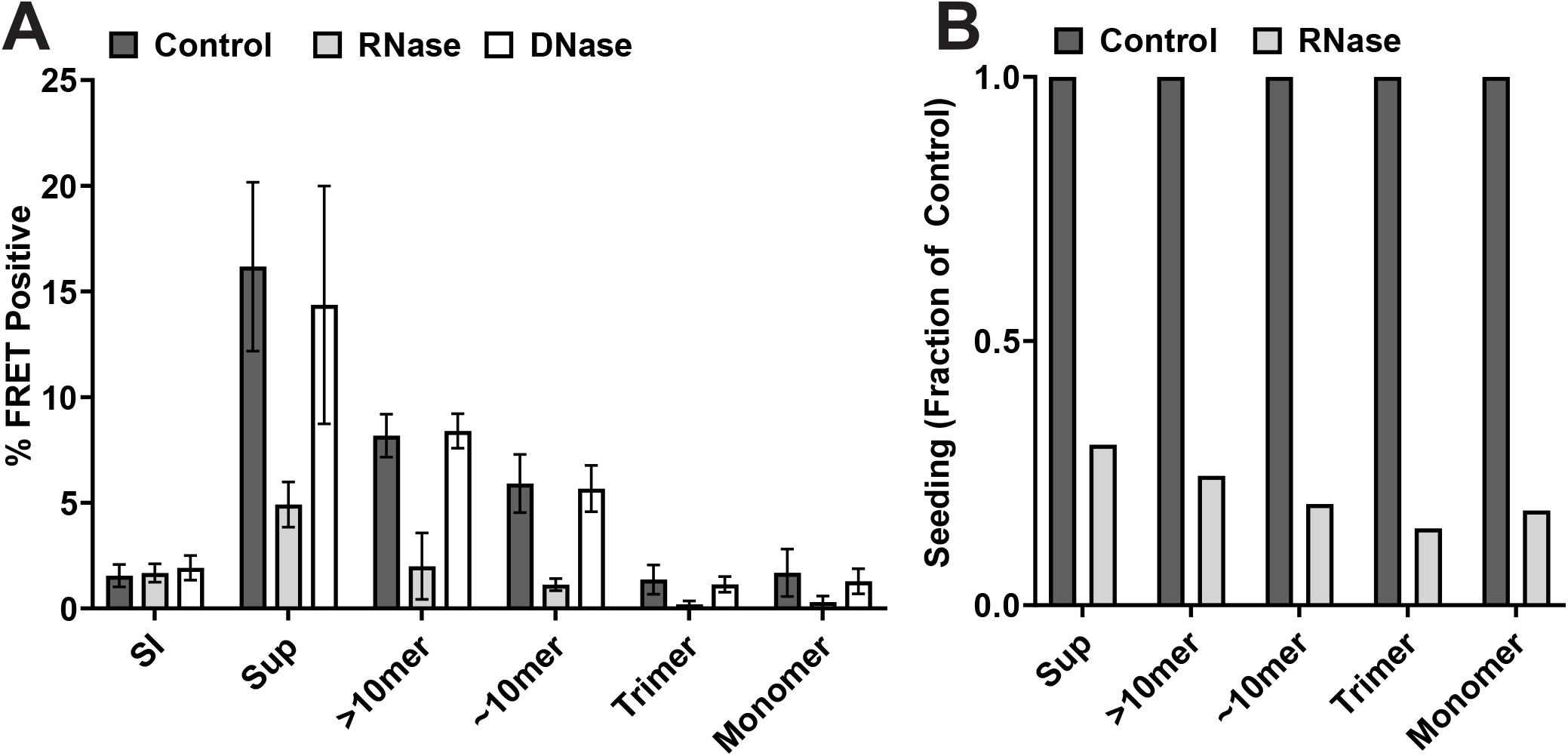
RNA stabilizes soluble seeds from AD brain. Homogenate from AD brain was fractionated in sarkosyl via ultra-centrifugation. The supernatant (sup) was further fractionated using SEC. (A) Resultant fractions were treated with RNase, DNase, or vehicle for 24hrs and seeded onto v2L biosensors. The percentage of FRET-positive cells was determined at 48hrs. Soluble AD seeds (Sup, >10mer, ˜10mer, trimer, monomer) were highly sensitive to RNase (All P<0.01 unpaired t-test as compared to control) but not DNase (P>0.45 unpaired t-test as compared to control). Sarkosyl Insoluble (SI) AD fibrils were unchanged in all conditions. Graphs represent data from two experimental replicates, each performed in technical triplicate. These experiments are representative of similar studies performed 12 times overall. Error bars = S.D. (B) Data from A normalized to untreated control seeding (% FRET Positive).

## Discussion

This study has examined the role of RNA in the development and maintenance of tau seeds *in vitro* and in AD. We initially found that tau bound multiple forms of single-stranded RNA with relatively high avidity, yet formation of seeds and fibrils depended on RNA sequence and size, with a maximally effective size >40nt. RNA was critical to maintaining induced seed integrity *in vitro*. While RNA from a variety of sources induced seed-competent tau, when this was used to transduce a human biosensor cell system, strain stability was far more robust when seeds had been produced by induction with human RNA. Finally, soluble tau seeding activity from AD brain was highly sensitive to RNase but not DNase, indicating a critical role for RNA in the maintenance of AD-derived seeds. While more studies are required, these observations suggest that certain RNAs might specifically trigger the formation and maintenance of unique tau assemblies, or strains, contributing to the diversity of tauopathies.

### RNA pathways, tau, and prions

Our finding that RNA modulates the conformation of tau assemblies builds on many prior reports that have linked tau and RNA-binding proteins (RBPs) in disease (reviewed in [30] and [31]. Tau has been reported to bind ribosome components, including rRNAs [32-37], and can affect ribosome biogenesis and mRNA translation ([35, 38-40]. Tau binds RBPs, such as TIA1 and SRRM2, that modulate RNA splicing and stress granule formation [20, 41-52]. Finally, tau directly binds RNA in inert [53] and seed-competent conformations [20]. Our findings suggest that beyond simple charge-based structural or scaffold effects, the tau:RNA complex may underlie physiological and pathophysiological processes based on conversion of tau between an inert and seed-competent state.

Like tau, the prion protein (PrP) may also use RNA as a cofactor to replicate unique structures (reviewed in [54]. RNA can facilitate PrP conversion *in vitro*, maintain infectivity in established prion strains, and act in a strain specific fashion [55-62]. Thus, our observations for tau may reflect a more general role for prions in regulating RNA, or vice versa.

### RNA binding vs. seed induction

We observed that tau binds single-stranded RNA independent of sequence, as avidities were similar for three homopolymeric RNAs. Given lack of knowledge about the binding mode of RNA to tau, we cannot refer to precise affinities, although the association/dissociation kinetics were most consistent with two RNA binding sites per tau molecule. Most studies involving tau and RNA have suggested non-specific associations based on simple charge interactions[16, 17, 63, 64]. One study suggested a preference for tRNA binding to tau overexpressed in cells[53]. By contrast, we observed that only some single-stranded forms of RNA induced tau assembly while other single-stranded and all double-stranded sequences did not. While ss DNA induced seeding, DNA exists largely as a double strand in cells, and ds DNA failed to induce tau fibrilization. This is consistent with prior reports [63, 65]. Other work has described polyU induction in tau of EPR signatures of in-register β-strands or increased ThT signal[65, 66]. In these studies, polydisperse RNA was incubated with repeat domain fragments (K18, K19), or mutated tau (P301L), which differs from the conditions used here, which were based exclusively on FL tau (2N4R) without disease-associated mutations or strong intrinsic self-assembly characteristics [67]. In summary, despite evidence that many forms of RNA bind tau and induce seeding and fibril formation, this work is the first to highlight the role of RNA as a specific inducer and stabilizer of tau seeds and strains that propagate in cells.

### RNA:tau and seed formation

While all ratios of polyA and tau generated seeds, 2:1 produced the most efficient seeding *in vitro*. This agreed with the 2:1 best fit of our binding data. The biphasic response of tau seeding to RNA concentration agreed with a prior report of optimal protein:inducer stoichiometry in the context of tau polymerization in presence of heparin and arachidonic acid[68]. RNAs did not induce tau seeding equally. We observed a minimum of 40 nt requirement to induce measurable seeding in sensitive biosensors.Despite evidence of relatively non-specific binding, which might have been mediated primarily by complementary charge interactions, RNA exerted a specific effect on tau seed formation, and especially on the induction of stable tau strains.

### RNA induction of strains

We observed that virtually all tau in ordered assemblies (whether derived from heparin, DNA, or RNA) seeded acutely into biosensor cells. However, only certain RNAs preferentially formed structures that propagated faithfully over multiple cell divisions, i.e. with strain characteristics. Importantly, in this study we did not isolate and characterize individual strains as we have done previously[4, 6, 8], which is highly labor-intensive.However we did characterize strains using the coarse-grain characteristic of inclusion morphology, and observed clear differences in strain composition based on the RNA inducer (Figure 3). We have previously observed that heparin-induced fibrils induce the formation of strains relatively inefficiently. In our original study, we observed that only about 1-2% DS1 cells transduced by heparin:tau stably propagated a strain[4].

This work puts these findings in a new light. Specifically, we found that the source of inducer used to create tau seeds had an enormous impact on the stability of the strains that subsequently propagate in cultured HEK293T cells. For example, when we used heparin or yeast RNA, we readily formed fibrils and seeds that induced aggregation acutely in cultured cells, but inclusions did not persist. By contrast, induction of tau with human RNA efficiently triggered the formation of more stable strains. Coupled with data which indicated that RNA played a critical role in stabilizing induced fibrils, and soluble AD seeds, these data suggest a role for RNA within the cell in the stabilization of strains. We hypothesize that strain stability in a cell will depend on the ability to bind specific classes, or even sequences of RNA.

According to this model (Figure 5), yeast RNAs do not form stable strains in HEK293T cells because the conformations of tau induced by these molecules do not encounter suitable sequences within the cell, and thus the strains are rendered unstable. By contrast, use of human RNA as an inducer *in vitro* creates a tau conformation competent to bind similar RNAs within the cell, and thus more stably propagates strains. These findings are consistent with the observation that RNA is associated with neurofibrillary tau deposits in human brain[18, 19]. An intriguing extension of this hypothesis is that rather than simply recruiting RNA in disease states, tau might play a regulatory role in RNA metabolism. This idea will require much more detailed study, but makes very specific predictions. First, RNA-dependent tau seeding activity should be found in normal conditions, and additionally specific strains will bind unique classes or sequences of RNA that may in turn be required for their formation and/or stable maintenance.

**Figure 5:**
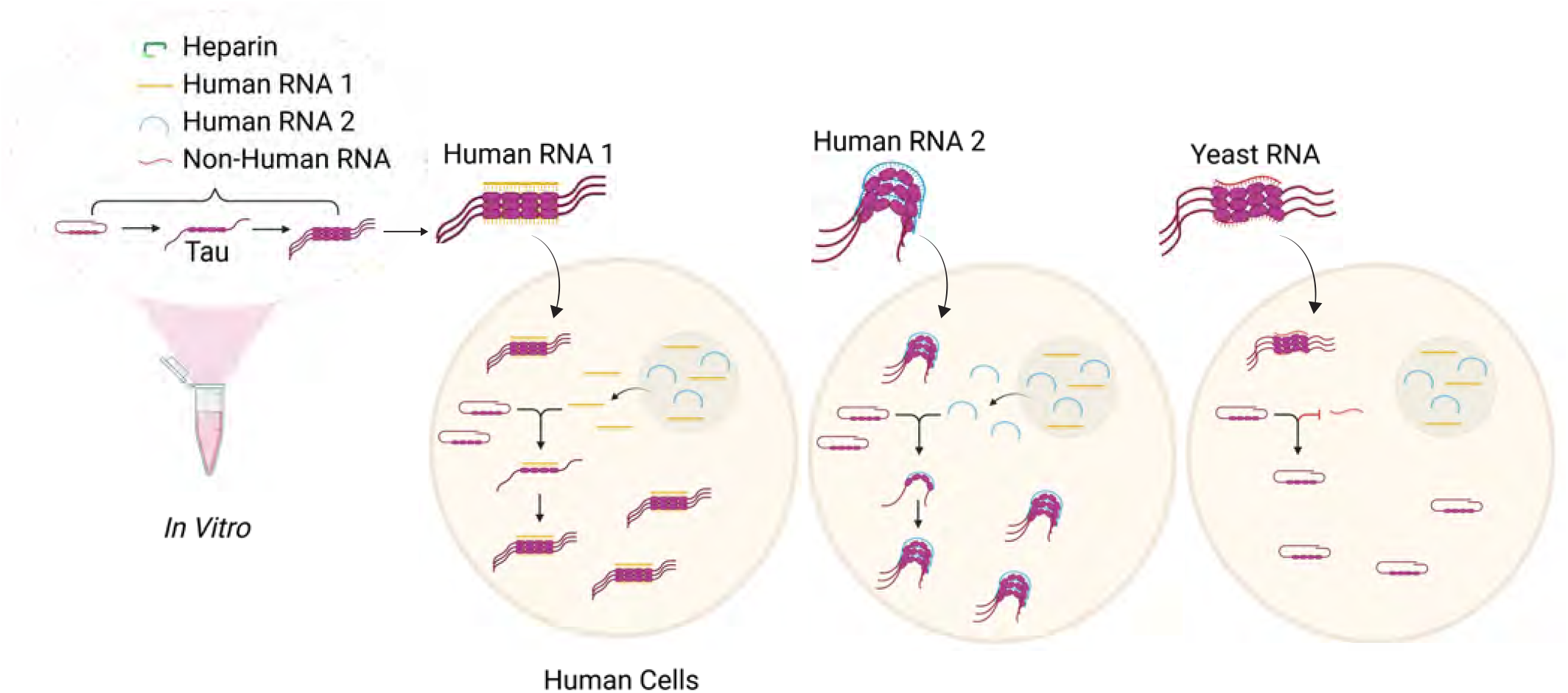
Model of RNA:tau seeding and strain propagation. Many inducers convert tau from inert to seed-competent to form distinct types of higher order assemblies *in vitro*. Unique RNAs are required to maintain distinct conformations, so when seeds are introduced into human cells, only those conformations that bind specific RNAs will propagate stable intracellular aggregates.

### RNA stabilizes soluble seeds from AD brain

Consistent with our observations that RNA stabilized tau seeds *in vitro*, we observed that RNase treatment profoundly diminished seeding activity all forms of soluble tau from AD brain, whereas DNase treatment had no effect. This implies that seed-competent tau detected in the biosensor cell system requires RNA to maintain its activity. Despite a documented association with RNA[18, 19], seeding activity of detergent-insoluble tau was unaffected by RNase treatment. This could be due to maturation of seeds into a fibrillar conformation that no longer requires RNA, or the contribution of other factors to fibril stability. Interestingly, it was recently reported that seeded soluble tau oligomers, and not fibrils, associate with RBPs, while fibrils are shuttled for autolysosomal degradation, suggesting unique physiological activity for oligomers [69]. While we cannot exclude RNA as simply a non-structural “co-factor “ required for seeding, taken together with *in vitro* studies, these data suggest a primary role for RNA in converting and maintaining tau in a seed-competent form in AD.

## Conclusion

These experiments have explored the role of RNA in the conversion of tau from an inert to a seed competent form, the formation of unique strains, the stability of pathogenic assemblies, including soluble seeds in AD. Whether specific RNAs bind AD seeds, and whether they govern initial transformation of tau from an inert to a seed-competent form demands further study, but could shed light on the mechanistic origins of tauopathy.

### Experimental Procedures

#### Cell culture

All cells were grown in Dulbecco’s Modified Eagle Medium (Gibco) supplemented with 10% fetal bovine serum (HyClone) and 1% Glutamax (Gibco). Cells were maintained at 37°C, 5% CO_2_, in a humidified incubator and routinely tested for mycoplasma (VenorGem, Sigma).

#### Tau expression and purification

We prepared full-length (FL) wild-type (WT) tau (2N4R) protein as previously described [25].

#### Nucleic acid purification

Total human RNA fractions were purified from HEK293T cells using Zymo Direct-zol RNA miniprep plus kits. All ssRNA and ssDNA sequences were synthesized by Sigma. Total yeast RNA (*Saccaromyces cerevisiae*) was purchased from Sigma. All nucleic acid preparations were stored in diethylpyrocarbonate (DEPC) treated H_2_O at −80°C.

#### Synthetic nucleic acid sequences

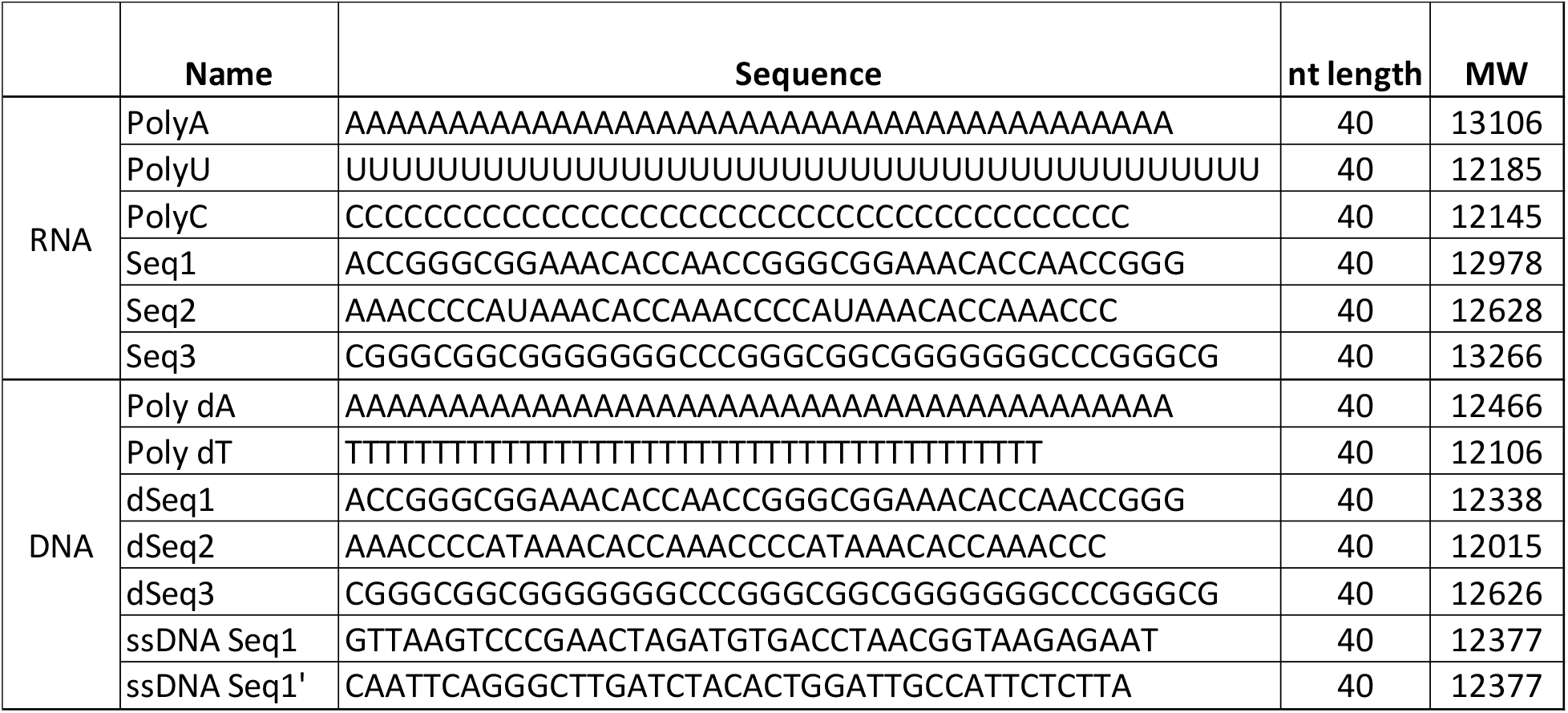

#### Fibrillization

FL WT tau monomer was filtered through a 100kDa molecular weight cut-off (MWCO) filter (Corning) as instructed by the manufacturer (15,000 x g for 15 minutes). Filtered protein was collected and protein concentration determined via DeNovix DS-11 Spectrophotometer. RNA or DNA was boiled for 7 minutes and snap-cooled on ice before addition to filtered monomer at determined mass ratio. All nucleic acid concentrations were quantified on a DeNovix DS-11 Spectrophotometer. Tau monomer was brought to a final concentration of 8µM using tau buffer (10mM HEPES, 100mM NaCl in Millipore H_2_O). DTT was added at a final concentration of 11% before adding inducer or buffer. Fibril mixtures were agitated at 350rpm using a Thermomixer (Eppendorf) set to 37°C for determined time points.

#### Seeding assay

Stable v2L biosensor cells[24], which express tau repeat domain containing the P301S mutation, fused to cerulean or mClover fluorescent proteins, were plated at a density of 30,000 cells/well in a 96-well plate. After 18 hr, at 60% confluency, cells were transduced with tau fibrils using Lipofectamine 2000 (Invitrogen). Cells were incubated with tau fibrils for 48 hours before harvesting for flow cytometry.

#### Flow cytometry

Transduced biosensor cells were harvested with 0.05% trypsin and fixed in 2% paraformaldehyde for 10 minutes before being resuspended in flow cytometry buffer (Hank’s Balanced Salt Solution supplemented with 1% fetal bovine serum and 1mM EDTA), and evaluated by flow cytometry (Fortessa, GE) as described previously [23, 24]. Three technical replicates were used for each data set. For each experiment, a minimum of 5000 single cells per replicate were analyzed. Data analysis was performed using FLOWJO v10 and GraphPad PRISM v9.

#### Strain-containing monoclonal generation

Stable cell lines expressing tau-repeat domain with P301L/V337M mutation fused to YFP (DS1) [4] were plated at a density of 25,000 cells/well in 96-well plates. Tau fibrils generated from RNA or DNA were transduced into cells using Lipofectamine 2000 and allowed to incubate for 72 hours. Cells were then trypsinized and resuspended in flow buffer before live-cell sorting. Aggregate-containing cells were identified by their bright YFP signal using a FACS Aria II SORP cell sorter, as previously described [8]. Cells were sorted individually into a 96-well plate and grown until confluency to derive monoclonal lines that propagated tau strains.

#### Aggregate maintenance assay

Monoclonal colonies containing aggregates grew in 96-well plates for two weeks before being scored for aggregates and amplified. Cell colonies were followed for two months, passaging into 24-well plates and 12-well plates before storage in liquid nitrogen.Colonies were scored as containing aggregates uniformly, or sectored, i.e. with loss of aggregates.

#### Fixed cell imaging

Stably passaged monoclonal cell lines were plated onto a Corning Special Optics 96-well plate and allowed to grow to ˜40% confluency before fixing in 4% paraformaldehyde and staining with DAPI. Cells were then imaged via confocal microscopy (IN CELL Analyzer 6000, GE). Each colony was imaged from 13 distributed points in each well, across 5 z-stacks. IN CELL software then compiled maximum intensity projections (MIPs) across z-stacks to create a single image from each point scanned. MIP files were coded and blindly scored for tau aggregate morphology before unblinding and quantification.

#### Transmission electron microscopy (TEM)

Fibrils (4μM) were deposited onto glow-discharged Formvar-coated 400-mesh copper grids for 30 sec, washed with distilled water, and then negatively stained with 2% uranyl acetate for 1 minute. Images were acquired on a Tecnai G^2^ spirit transmission electron microscope (FEI, Hillsboro, OR), serial number: D1067, equipped with LaB_6_ source at 120kV using a Gatan ultrascan CCD camera.

#### Biolayer interferometry

Second Generation Amine Reactive Biosensors (AR2G, ForteBio) were incubated with EDC and Sulfo-NHS to activate carboxylic acids. FL WT tau was diluted in acetate (pH 6) to charge primary amines and allow immobilization on biosensors. After quenching biosensors with ethanolamine, tau coated sensors were dipped into serial 3-fold dilutions (1/2 log) of RNA to determine association and disassociation kinetics. Data from wells containing no RNA, and biosensors containing no tau were background subtracted. Data were analyzed using Octet Data Analysis software using y-axis alignment, Savitz-Golay filtering, and inter-step correction.

#### Enzyme treatment

Pre-formed tau fibrils were incubated with a mixture of RNase A (3.2 mg/mL, Thermo) and RNase T1 (316 U/μL, Thermo), DNase I (633 U/μL, NEB), Heparinase I (3797U/mL) or Millipore H_2_O (Control) and were agitated at 750 RPM in a Thermomixer (Eppendorf) set to 37°C for 24 hr.

#### Human brain homogenization

0.5g sections of frontal lobe from AD brain were dounce homogenized in PHF buffer (10mM Tris-HCl (pH 7.4), 0.8M NaCl, 1mM EGTA, and 10% sucrose) [70], and sarkosyl insoluble fractions prepared as previously described [71] using ultra-centrifugation at 186,000 x g for 1 hour at 4°C to separate pellet from supernatant. Supernatant was flash frozen and stored at −80°C for subsequent size fractionation. Pellets were washed with PBS for 30 minutes at 186,000 x g, 4°C and resuspended in 50mM Tris-HCL (pH 7.5) before flash freezing and storing at −80°C.

#### Size exclusion chromatography (SEC)

AD ultra-centrifuged supernatant was loaded onto a Superdex 200 Increase 10/300 GL Column (GE Healthcare). SEC fractions were eluted in PHF buffer. Fractions were concentrated using 2000kD MWCO Filters (Vivacon), evaluated for protein concentration using A215 spectra, and frozen at −80°C.

#### Data availability

All data generated and analyzed during this study are included in this article.

## Supporting information

This article contains supporting information.

## Acknowledgements

The authors would like to thank the members of the Center for Alzheimer’s and Neurodegenerative Diseases for helpful critiques and discussions of this work, particularly: Michael Abrams, Jordan Finnell, Hilda Mirbaha, Anuja Modi, Apurwa Sharma, William Russ, David Sanders, and DaNae Woodard. This work was supported by the Chan Zuckerberg Initiative and NIH/NIA 1R01AG059689. We appreciate the assistance of the Moody Foundation Flow Cytometry Core Facility and UT Southwestern Electron Microscopy Core, especially Phoebe Doss and Becca Jackson.

## Author contributions

MID and ANZ designed research; ANZ, OMK, JEC, CLW performed research and/or contributed reagents; MID and AZZ wrote the manuscript.

## Funding and additional information

This work was supported by the Chan Zuckerberg Initiative and NIH/NIA 1R01AG059689. The content is solely the responsibility of the authors and does not necessarily represent the official views of the National Institutes of Health.

## Supporting Information

**supplemental Figure 1:**
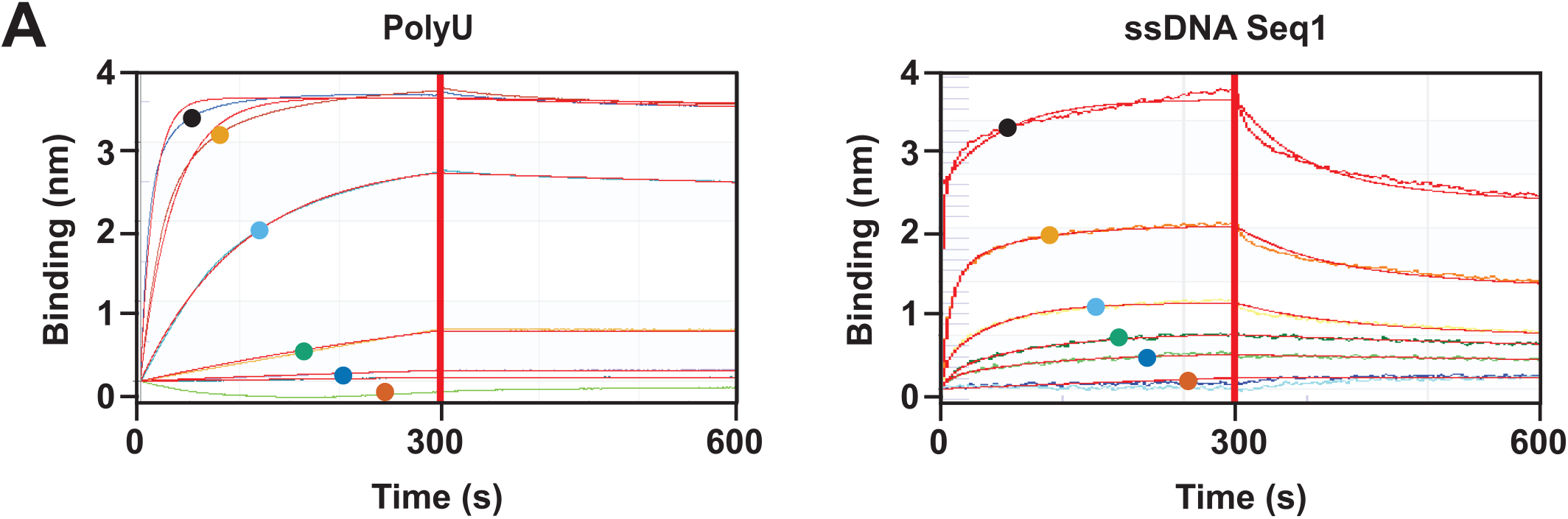
Additional binding curves. (A) BLI with tau immobilized onto AR2G biosensors and exposed to increasing concentrations of PolyU RNA, or ssDNA Seq1 (41μM (•), 13μM 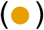 4μM 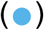, 1.3μM 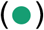, 0.4μM 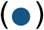, 0.13μM 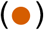. Figure shows representative data of PolyU RNA or ssDNA Seq1 binding to BLI AR2G sensors loaded with tau. K_DAPP_ calculated as mean of three independent experiments. Binding (nm) refers to the perturbation in reflected light from the biosensor. Red lines represent curves fit to primary data. (See also Figure 1A; Table 1)

**supplemental Figure 2:**
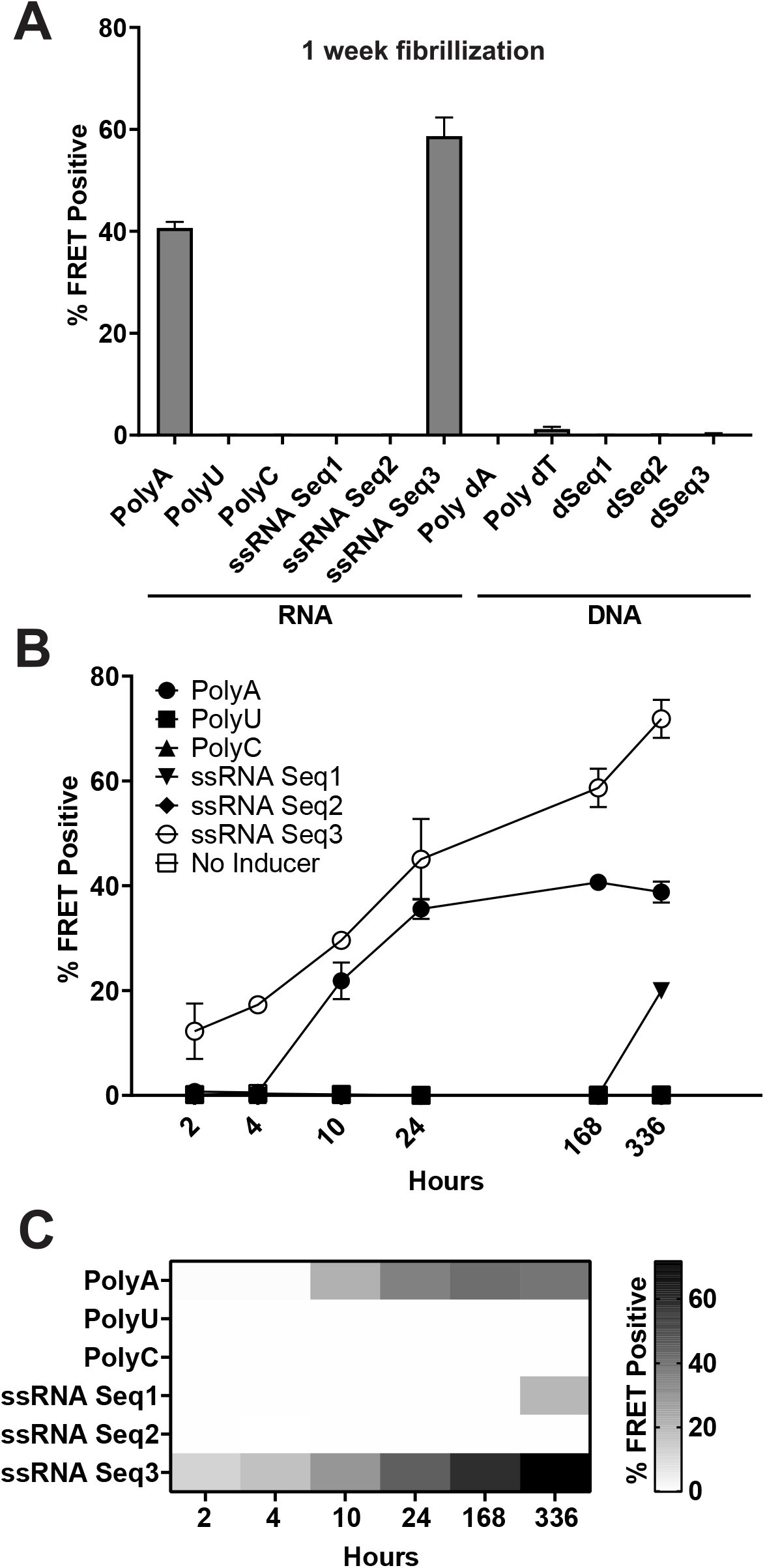
Time course of nucleic acid seeding. (A) Tau monomer was incubated 1 week with nucleic acid before transduction of v2L biosensors. We quantified intracellular aggregation as % FRET positive via flow cytometry. Specific nucleic acids induced seed-competency in tau (Poly A, ssRNA Seq3) while others did not (PolyU, PolyC, ssRNA Seq1-2, PolydA, PolydT, dSeq1-3). (B) Time course of tau incubation with different nucleic acids over time. (C) Heatmap of data in B. All conditions quantified in technical triplicate. Error bars = S.D.

**supplemental Figure 3:**
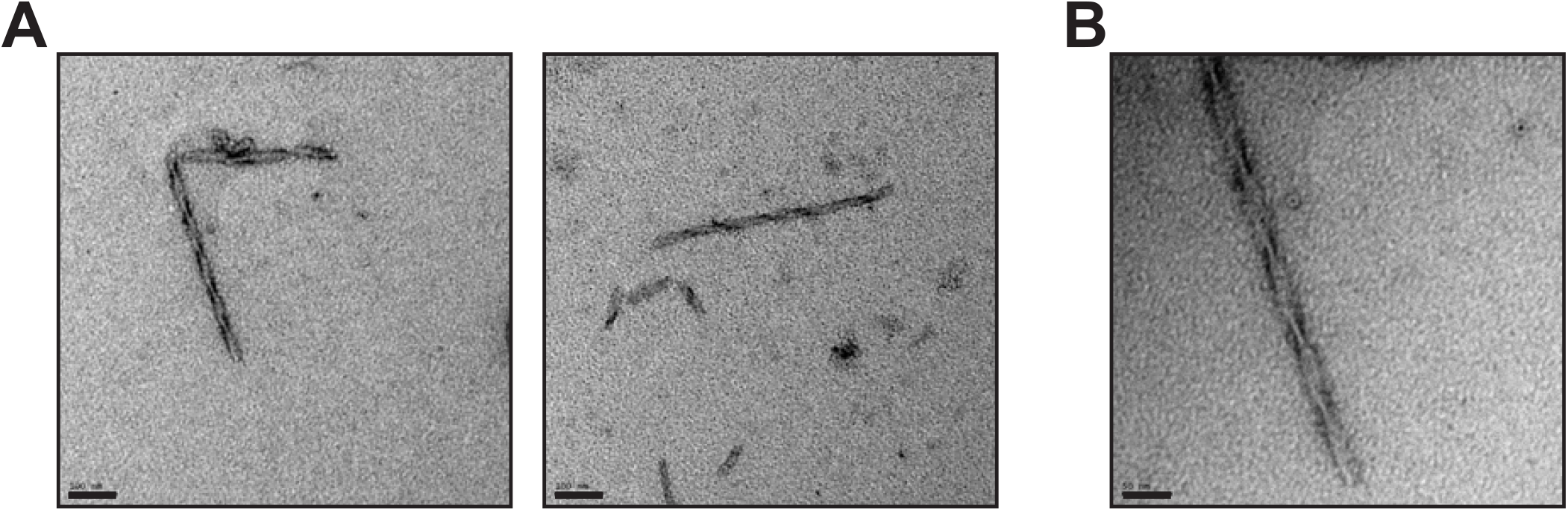
SI extraction of tau fibrils from AD brain. After dounce homogenizing AD brain, insoluble fibrils were fractionated using sarkosyl and ultra-centrifugation. Fibrils were negative stained and imaged with TEM. (A) Fibrils, scale bar = 100nM, (B) scale bar = 50nM.

**supplemental Figure 4:**
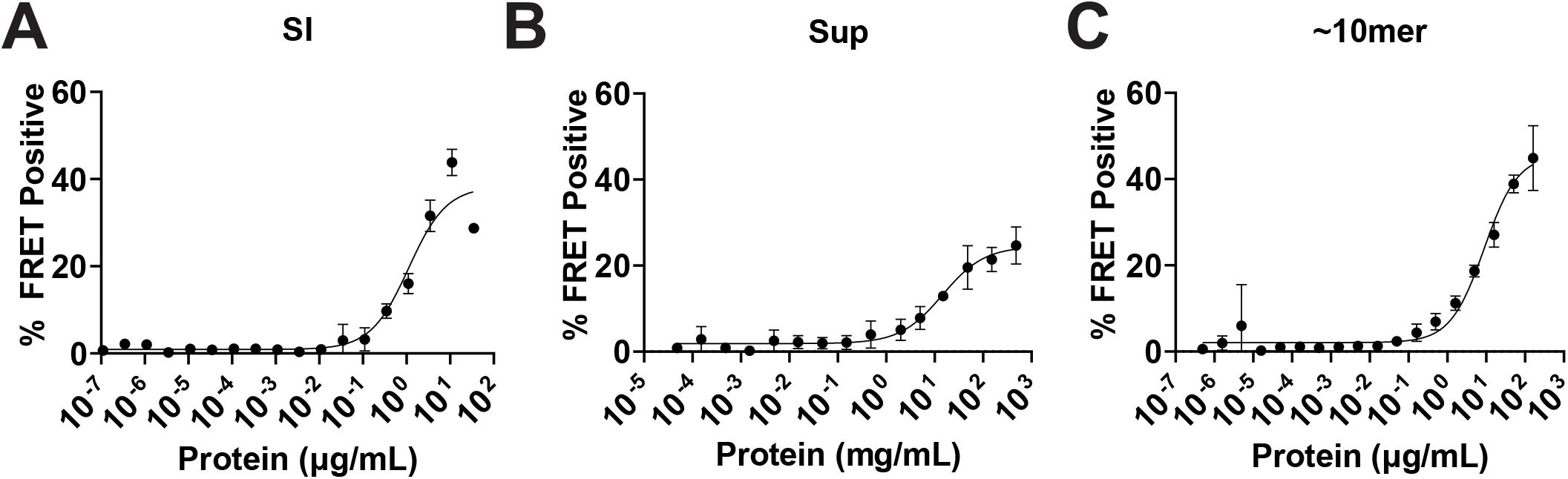
Titration of AD seed fractions on v2L cells. AD homogenate was fractionated in sarkosyl with ultra-centrifugation. Supernatant was further fractionated using SEC. Increasing protein amounts of (A) sarkosyl insoluble (SI), (B) supernatant, or (C) ˜10mer were transduced into v2L cells and intracellular seeding was quantified after 48 hr. Data were fit with nonlinear least squares regression curves to determine linear range of protein to use for subsequent nuclease experiments. All data points represent technical triplicates. Error bars = S.D.

## References

1. Lee, V.M., M. Goedert, and J.Q. Trojanowski, Neurodegenerative tauopathies. Annu Rev Neurosci, 2001. 24: p. 1121–59.

2. Frost, B., R.L. Jacks, and M.I. Diamond, Propagation of tau misfolding from the outside to the inside of a cell. J Biol Chem, 2009. 284(19): p. 12845–52.

3. Clavaguera, F., et al., Transmission and spreading of tauopathy in transgenic mouse brain. Nat Cell Biol, 2009. 11(7): p. 909–13.

4. Sanders, D.W., et al., Distinct tau prion strains propagate in cells and mice and define different tauopathies. Neuron, 2014. 82(6): p. 1271–88.

5. Calafate, S., et al., Synaptic Contacts Enhance Cell-to-Cell Tau Pathology Propagation. Cell Rep, 2015. 11(8): p. 1176–83.

6. Kaufman, S.K., et al., Tau Prion Strains Dictate Patterns of Cell Pathology, Progression Rate, and Regional Vulnerability In Vivo. Neuron, 2016. 92(4): p. 796–812.

7. Clavaguera, F., et al., Brain homogenates from human tauopathies induce tau inclusions in mouse brain. Proc Natl Acad Sci U S A, 2013. 110(23): p. 9535–40.

8. Sharma, A.M., et al., Tau monomer encodes strains. Elife, 2018. 7.

9. Wesseling, H., et al., Tau PTM Profiles Identify Patient Heterogeneity and Stages of Alzheimer’s Disease. Cell, 2020. 183(6): p. 1699-1713.e13.

10. Mirbaha, H., et al., Seed-competent tau monomer initiates pathology in PS19 tauopathy mice. bioRxiv, 2022: p. 2022.01.03.474806.

11. Friedhoff, P., et al., Rapid assembly of Alzheimer-like paired helical filaments from microtubule-associated protein tau monitored by fluorescence in solution. Biochemistry, 1998. 37(28): p. 10223–30.

12. Goedert, M., Jakes, R., Spillantini MG, Hasegawa M, Smith MJ, Crowther RA, Assembly of microtubule-associated protein tau into Alzheimer-like filaments induced by sulphated glycosaminoglycans. Nature, 1996. 383: p. 550–553.

13. Pérez, M., et al., Polymerization of tau into filaments in the presence of heparin: the minimal sequence required for tau-tau interaction. J Neurochem, 1996. 67(3): p. 1183–90.

14. Wilson, D.M. and L.I. Binder, Free fatty acids stimulate the polymerization of tau and amyloid beta peptides. In vitro evidence for a common effector of pathogenesis in Alzheimer’s disease. Am J Pathol, 1997. 150(6): p. 2181–95.

15. Chirita, C.N., M. Necula, and J. Kuret, Anionic micelles and vesicles induce tau fibrillization in vitro. J Biol Chem, 2003. 278(28): p. 25644–50.

16. Kampers, T., et al., RNA stimulates aggregation of microtubule-associated protein tau into Alzheimer-like paired helical filaments. FEBS Lett, 1996. 399(3): p. 344–9.

17. Bryan, J.B., B.W. Nagle, and K.H. Doenges, Inhibition of tubulin assembly by RNA and other polyanions: evidence for a required protein. Proc Natl Acad Sci U S A, 1975. 72(9): p. 3570–4.

18. Ginsberg, S.D., et al., Sequestration of RNA in Alzheimer’s disease neurofibrillary tangles and senile plaques. Ann Neurol, 1997. 41(2): p. 200–9.

19. Ginsberg, S.D., et al., RNA sequestration to pathological lesions of neurodegenerative diseases. Acta Neuropathol, 1998. 96(5): p. 487–94.

20. Lester, E., et al., Tau aggregates are RNA-protein assemblies that mislocalize multiple nuclear speckle components. Neuron, 2021. 109(10): p. 1675-1691.e9.

21. Gustke, N., et al., Domains of tau protein and interactions with microtubules. Biochemistry, 1994. 33(32): p. 9511–22.

22. Mukrasch, M.D., et al., The “jaws” of the tau-microtubule interaction. J Biol Chem, 2007. 282(16): p. 12230–9.

23. Holmes, B.B., et al., Proteopathic tau seeding predicts tauopathy in vivo. Proc Natl Acad Sci U S A, 2014. 111(41): p. E4376–85.

24. Hitt, B.D., et al., Ultrasensitive tau biosensor cells detect no seeding in Alzheimer’s disease CSF. Acta Neuropathol Commun, 2021. 9(1): p. 99.

25. Frost, B., et al., Conformational diversity of wild-type Tau fibrils specified by templated conformation change. J Biol Chem, 2009. 284(6): p. 3546–51.

26. Hartl, F.U., Protein Misfolding Diseases. Annual Review of Biochemistry, 2017. 86(1): p. 21–26.

27. Mirbaha, H., et al., Inert and seed-competent tau monomers suggest structural origins of aggregation. Elife, 2018. 7.

28. Hou, Z., et al., Biophysical properties of a tau seed. Sci Rep, 2021. 11(1): p. 13602.

29. Holtzman, D.M., J.C. Morris, and A.M. Goate, Alzheimer’s disease: the challenge of the second century. Sci Transl Med, 2011. 3(77): p. 77sr1.

30. Koren, S.A., S. Galvis-Escobar, and J.F. Abisambra, Tau-mediated dysregulation of RNA: Evidence for a common molecular mechanism of toxicity in frontotemporal dementia and other tauopathies. Neurobiology of Disease, 2020. 141: p. 104939.

31. Rybak-Wolf, A. and M. Plass, RNA Dynamics in Alzheimer’s Disease. Molecules, 2021. 26(17).

32. Gunawardana, C.G., et al., The Human Tau Interactome: Binding to the Ribonucleoproteome, and Impaired Binding of the Proline-to-Leucine Mutant at Position 301 (P301L) to Chaperones and the Proteasome. Molecular & cellular proteomics : MCP, 2015. 14(11): p. 3000–3014.

33. Meier, S., et al., Pathological Tau Promotes Neuronal Damage by Impairing Ribosomal Function and Decreasing Protein Synthesis. J Neurosci, 2016. 36(3): p. 1001–7.

34. Meier, S., et al., Identification of Novel Tau Interactions with Endoplasmic Reticulum Proteins in Alzheimer’s Disease Brain. J Alzheimers Dis, 2015. 48(3): p. 687–702.

35. Koren, S.A., et al., Tau drives translational selectivity by interacting with ribosomal proteins. Acta Neuropathol, 2019. 137(4): p. 571–583.

36. Banerjee, S., et al., Tau protein-induced sequestration of the eukaryotic ribosome: Implications in neurodegenerative disease. Sci Rep, 2020. 10(1): p. 5225.

37. Jessus, C., et al., In vitro inhibition of tubulin assembly by a ribonucleoprotein complex associated with the free ribosome fraction isolated from Xenopus laevis oocytes: effect at the level of microtubule-associated proteins. Cell Differ, 1984. 14(3): p. 179–87.

38. Evans, H.T., et al., Altered ribosomal function and protein synthesis caused by tau. Acta Neuropathol Commun, 2021. 9(1): p. 110.

39. Montalbano, M., et al., Tau Modulates mRNA Transcription, Alternative Polyadenylation Profiles of hnRNPs, Chromatin Remodeling and Spliceosome Complexes. Frontiers in Molecular Neuroscience, 2021. 14(302).

40. Evans, H.T., et al., Decreased synthesis of ribosomal proteins in tauopathy revealed by non-canonical amino acid labelling. Embo j, 2019. 38(13): p. e101174.

41. Wolozin, B., Regulated protein aggregation: stress granules and neurodegeneration. Mol Neurodegener, 2012. 7: p. 56.

42. Apicco, D.J., et al., Reducing the RNA binding protein TIA1 protects against tau-mediated neurodegeneration in vivo. Nat Neurosci, 2018. 21(1): p. 72–80.

43. Hsieh, Y.C., et al., Tau-Mediated Disruption of the Spliceosome Triggers Cryptic RNA Splicing and Neurodegeneration in Alzheimer’s Disease. Cell Rep, 2019. 29(2): p. 301-316.e10.

44. Vanderweyde, T., et al., Interaction of tau with the RNA-Binding Protein TIA1 Regulates tau Pathophysiology and Toxicity. Cell Rep, 2016. 15(7): p. 1455–1466.

45. Bishof, I., et al., RNA-binding proteins with basic-acidic dipeptide (BAD) domains self-assemble and aggregate in Alzheimer’s disease. J Biol Chem, 2018. 293(28): p. 11047–11066.

46. Hales, C.M., et al., U1 small nuclear ribonucleoproteins (snRNPs) aggregate in Alzheimer’s disease due to autosomal dominant genetic mutations and trisomy 21. Mol Neurodegener, 2014. 9: p. 15.

47. Hales, C.M., et al., Changes in the detergent-insoluble brain proteome linked to amyloid and tau in Alzheimer’s Disease progression. Proteomics, 2016. 16(23): p. 3042–3053.

48. Bai, B., et al., U1 small nuclear ribonucleoprotein complex and RNA splicing alterations in Alzheimer’s disease. Proceedings of the National Academy of Sciences, 2013. 110(41): p. 16562.

49. Maziuk, B.F., et al., RNA binding proteins co-localize with small tau inclusions in tauopathy. Acta Neuropathologica Communications, 2018. 6(1): p. 71.

50. Jiang, L., et al., TIA1 regulates the generation and response to toxic tau oligomers. Acta Neuropathol, 2019. 137(2): p. 259–277.

51. Johnson, E.C.B., et al., Deep proteomic network analysis of Alzheimer’s disease brain reveals alterations in RNA binding proteins and RNA splicing associated with disease. Molecular Neurodegeneration, 2018. 13(1): p. 52.

52. Jiang, L., et al., Interaction of tau with HNRNPA2B1 and N(6)-methyladenosine RNA mediates the progression of tauopathy. Mol Cell, 2021. 81(20): p. 4209-4227.e12.

53. Zhang, X., et al., RNA stores tau reversibly in complex coacervates. PLoS Biol, 2017. 15(7): p. e2002183.

54. Weissmann, C., A ‘unified theory’ of prion propagation. Nature, 1991. 352(6337): p. 679–83.

55. Burke, C.M., et al., Cofactor and glycosylation preferences for in vitro prion conversion are predominantly determined by strain conformation. PLoS Pathog, 2020. 16(4): p. e1008495.

56. Deleault, N.R., et al., Protease-resistant prion protein amplification reconstituted with partially purified substrates and synthetic polyanions. J Biol Chem, 2005. 280(29): p. 26873–9.

57. Deleault, N.R., et al., Formation of native prions from minimal components in vitro. Proc Natl Acad Sci U S A, 2007. 104(23): p. 9741–6.

58. Deleault, N.R., R.W. Lucassen, and S. Supattapone, RNA molecules stimulate prion protein conversion. Nature, 2003. 425(6959): p. 717–20.

59. Adler, V., et al., Small, highly structured RNAs participate in the conversion of human recombinant PrP(Sen) to PrP(Res) in vitro. J Mol Biol, 2003. 332(1): p. 47–57.

60. Saá, P., et al., Strain-specific role of RNAs in prion replication. J Virol, 2012. 86(19): p. 10494–504.

61. Gonzalez-Montalban, N., et al., Changes in prion replication environment cause prion strain mutation. Faseb j, 2013. 27(9): p. 3702–10.

62. Katorcha, E., et al., Prion replication environment defines the fate of prion strain adaptation. PLoS Pathog, 2018. 14(6): p. e1007093.

63. Hasegawa, M., et al., Alzheimer-like changes in microtubule-associated protein Tau induced by sulfated glycosaminoglycans. Inhibition of microtubule binding, stimulation of phosphorylation, and filament assembly depend on the degree of sulfation. J Biol Chem, 1997. 272(52): p. 33118–24.

64. Wang, X., et al., The proline-rich domain and the microtubule binding domain of protein tau acting as RNA binding domains. Protein Pept Lett, 2006. 13(7): p. 679–85.

65. Dinkel, P.D., et al., RNA Binds to Tau Fibrils and Sustains Template-Assisted Growth. Biochemistry, 2015. 54(30): p. 4731–40.

66. Fichou, Y., et al., Cofactors are essential constituents of stable and seeding-active tau fibrils. Proc Natl Acad Sci U S A, 2018. 115(52): p. 13234–13239.

67. Barghorn, S. and E. Mandelkow, Toward a unified scheme for the aggregation of tau into Alzheimer paired helical filaments. Biochemistry, 2002. 41(50): p. 14885–96.

68. Carlson, S.W., et al., A Complex Mechanism for Inducer Mediated Tau Polymerization. Biochemistry, 2007. 46(30): p. 8838–8849.

69. Jiang, L., et al., Tau Oligomers and Fibrils Exhibit Differential Patterns of Seeding and Association With RNA Binding Proteins. Frontiers in Neurology, 2020. 11(1183).

70. Goedert, M., et al., Tau proteins of alzheimer paired helical filaments: Abnormal phosphorylation of all six brain isoforms. Neuron, 1992. 8(1): p. 159–168.

71. Mirbaha, H., et al., Tau Trimers Are the Minimal Propagation Unit Spontaneously Internalized to Seed Intracellular Aggregation. J Biol Chem, 2015. 290(24): p. 14893–903.

